# Regulatory element modules as universal features for single-cell chromatin analysis

**DOI:** 10.64898/2025.12.10.692786

**Authors:** Chrysania Lim, Javen Tan Yih Ruay, Tim Stuart

## Abstract

Single-cell chromatin accessibility data provide important insights into the activity of DNA regulatory elements in health and disease. However, the analysis of these data is made challenging by the lack of a common set of features for use in downstream analysis. This results in individual studies quantifying dataset-specific peak regions that cannot be directly compared to other studies. To address this challenge, we developed a comprehensive set of DNA regulatory element modules (REMO) for the human genome. Here we show how REMO can be applied to single-cell chromatin data to better separate cell states in a low-dimensional space compared to peak matrix quantification, greatly improve the scalability of dimension reduction steps, and enable automated annotation of cell types. This is accompanied by new memory-efficient and scalable software for the quantification of single-cell chromatin accessibility data.

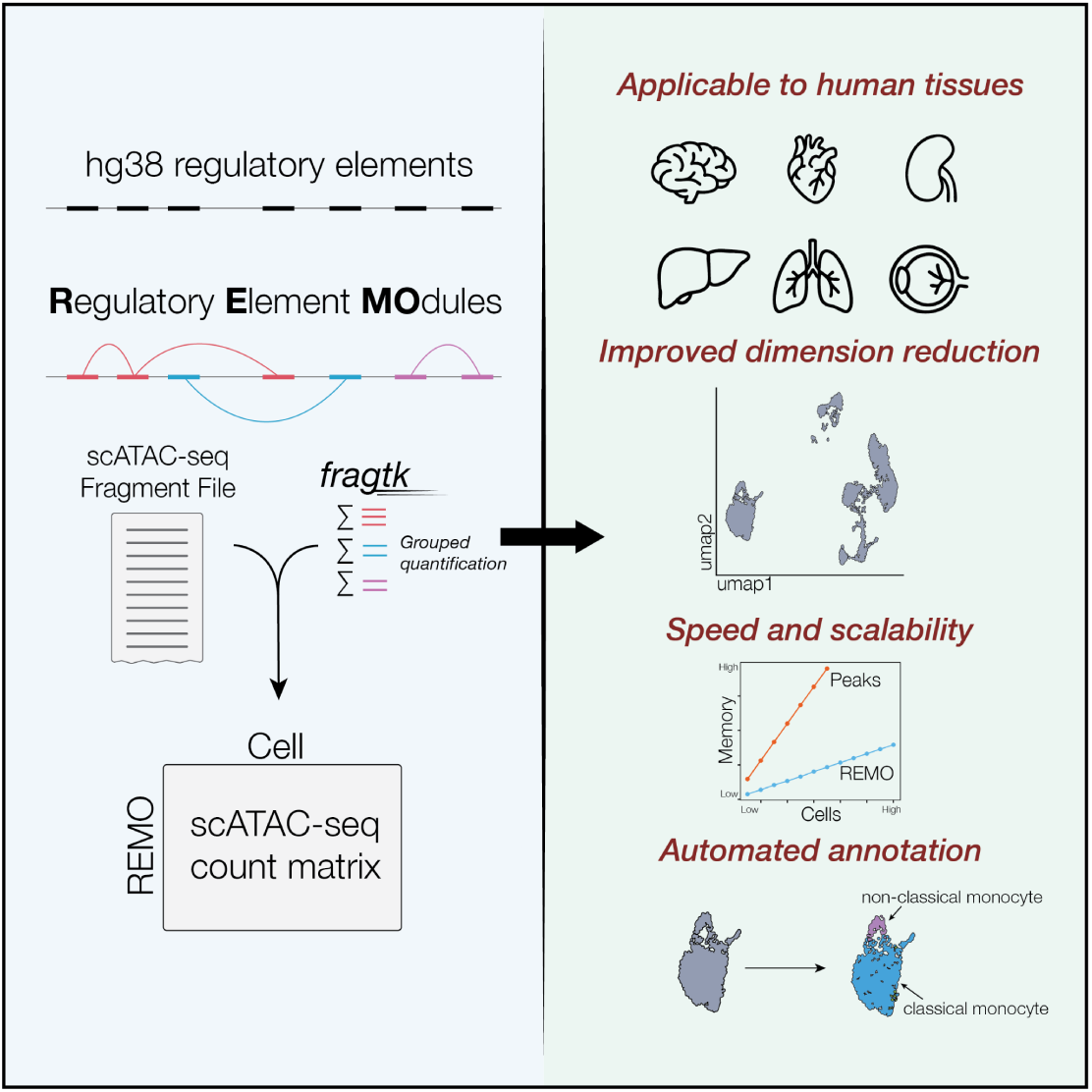

## Introduction

Single-cell chromatin profiling methods have enabled important insights into the molecular states of DNA regulatory elements across a broad diversity of cell types. These insights are critical to understanding how genes are regulated, how these regulatory processes are altered under different disease conditions, and how regulatory elements are impacted by genetic variation. Experimental methods capable of generating single-cell chromatin accessibility data, such as scATAC-seq, are rapidly improving and it is now possible to generate high-quality data at scale ^1,2^. Massive-scale generation of scATAC-seq data is poised to substantially improve our understanding of gene regulation. However, the analysis of such data faces a number of major challenges. Foremost among these is a lack of common features that can be re-used across multiple experiments, analogous to a canonical set of genes used in the analysis of single-cell transcriptomic data.

Current approaches for the analysis of scATAC-seq data typically begin by defining a set of features (peaks or genomic windows) for quantification and further analysis. Defining these features *de novo* from the data results in several important problems. First, the features are unique to a particular dataset, preventing any straightforward comparison with other existing datasets. This becomes an increasingly important challenge as the number of published datasets continues to grow and there is an increasing need to analyze new data in the context of other existing datasets, or to create large-scale integrated atlas datasets incorporating published data from many independent sources. Second, the number of features identified is large, typically comprising over 100,000 genomic regions, and creates challenges in downstream processing. Performing dimension reduction steps is computationally challenging due to the size and sparsity of the count matrix. Finally, peak regions alone are difficult to interpret, and the annotation of cell types in scATAC-seq datasets remains a challenging issue due to a lack of annotated reference datasets utilizing a standardized set of genomic features.

To address these challenges we developed a set of universal features for human scATAC-seq analysis. We identified groups of highly similar regulatory elements in the human genome and aggregated these together as sets of regulatory <u>e</u>lement <u>mo</u>dules (REMO). Here, we demonstrate how REMO improves the analysis of human scATAC-seq data by improving the separation of cell states in low dimensions, scalability, and automated annotation of cell clusters. We provide user-friendly computational tools to enable broad adoption of REMO, including new memory-efficient and scalable data quantification software.

## Results

### Identification of regulatory element modules for the human genome

Previous efforts have identified comprehensive collections of regulatory elements in the human genome. This includes efforts from the ENCODE consortium in defining a set of candidate cis-regulatory elements (CREs) ^3^, and a set of consensus ATAC-seq peaks identified across a large collection of human tissues (cPeaks) ^4^. These collections include over 1 million CREs each. While these represent comprehensive catalogues of CREs, their practical application in single-cell analysis is hampered by their sheer number. Quantifying these features in scATAC-seq data results in extremely large and sparse count matrices that are computationally difficult to process.

We reasoned that many CREs are highly redundant, displaying similar patterns of DNA accessibility across cell types. The DNA accessibility of correlated CREs may therefore be summed for a cell without loss of information. This would substantially reduce the total number of features and reduce the sparsity of each measurement, and still enable such features to be universally applicable to different datasets. Extensive epigenomic data are now available for the human genome through ENCODE and other large-scale projects ^3^. We aimed to leverage these collections of epigenomic data to group similar regulatory elements together into regulatory element modules.

We first assembled a comprehensive collection of CREs for the human hg38 genome assembly by combining CREs in ENCODE and cPeaks, resulting in a set of 1,520,441 CREs. We grouped these CREs into modules based on their patterns of biochemical marks as measured by ENCODE histone and transcription factor ChIP-seq data across 4,157 experiments, and a co-accessibility-by-contact score (Figure 1A). To estimate the co-accessibility between CREs we assembled a large collection of scATAC-seq data from published studies, spanning 551 cell clusters from a variety of tissues. The co-accessibility between CREs was defined as the Pearson correlation in DNA accessibility across all pseudobulk cell clusters in this dataset. We developed a contact probability score by combining contact probability estimates from a collection of deeply sequenced Hi-C data ^3,5^ with a genomic distance-based kernel weight. By computing co-accessibility-by-contact between each CRE pair we constructed a graph where each node represented a CRE and the edges represented the co-accessibility-by-contact score between CREs. Graph-based clustering methods were applied to each chromosome to produce a set of 340,069 regulatory element modules, with each CRE being uniquely assigned to one module (Figure 1A).

**Figure 1:**
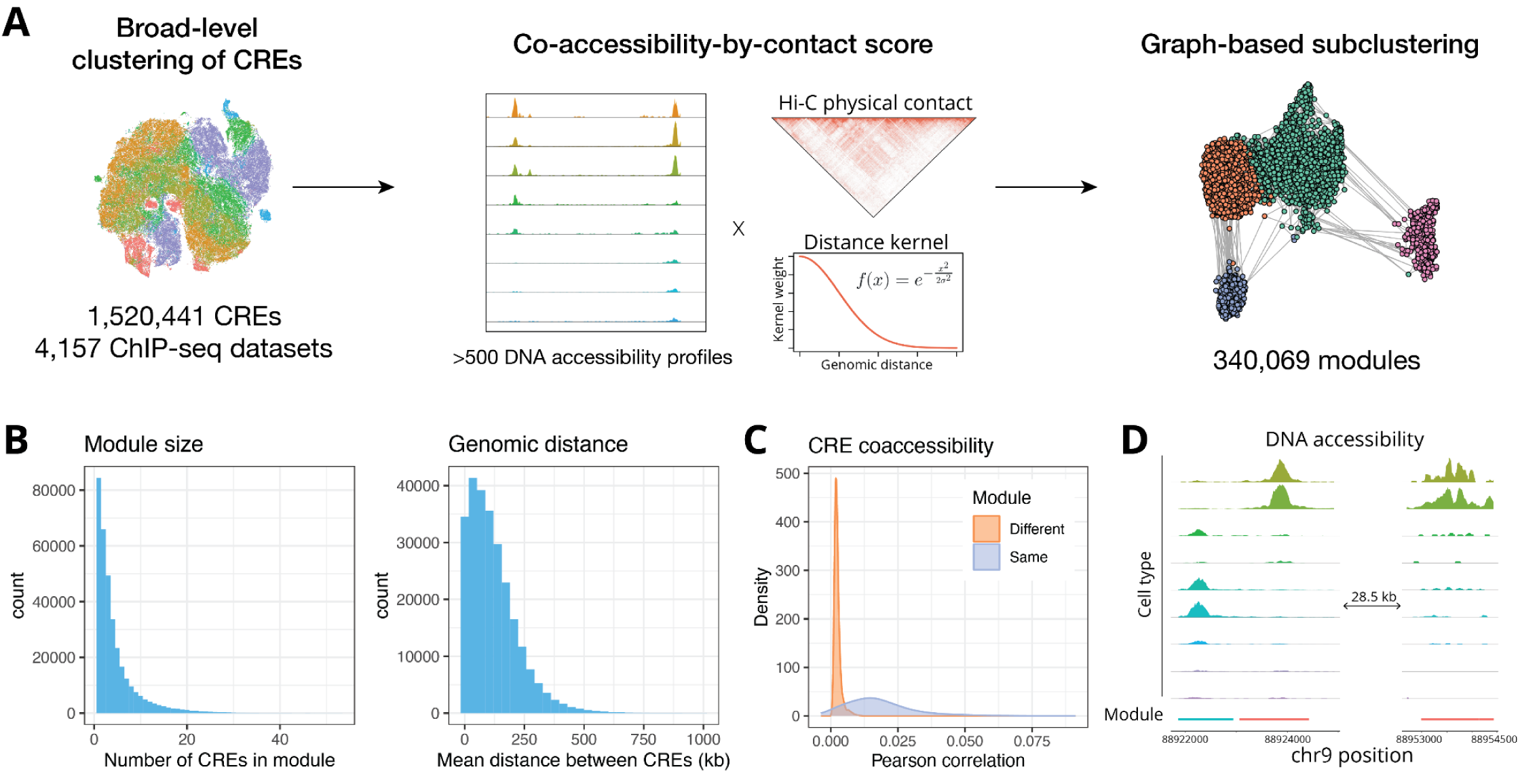
Identification of co-accessible regulatory element modules. **(A)** Schematic representation of the workflow used to identify regulatory element modules. CREs were first grouped into large sets of broadly similar elements for each chromosome using a collection of histone modification and transcription factor ChIP-seq data from the ENCODE consortium. Within each broad-level cluster, a CRE graph was constructed by computing pairwise co-accessibility-by-contact scores, using a large collection of pseudobulk cell cluster DNA accessibility profiles from scATAC-seq datasets, deeply sequenced bulk-cell Hi-C data, and a distance-based kernel weight. **(B)** Distribution of the number of CREs within each REMO module, and the pairwise genomic distance between CREs within the same module. **(C)** Distribution of Pearson correlation coefficient values between individual CREs in single-cell data, for CREs within the same module (blue) or in different modules (orange). **(D)** Representative genome browser visualization of CREs and their module grouping. Two CREs located adjacent to one another but displaying different cell type accessibility patterns are placed into different REMO modules, while another CRE further away in the genome (28,500 bp away) with similar cell type accessibility patterns is placed together in the same module. CREs are depicted using a colored bar showing their start and end coordinates, with the bar color corresponding to the assigned REMO module.

Overall, each module contained approximately 4 CREs (mean 4.4, sd 4.5), with the mean genomic distance between CREs within a module 120 kb (Figure 1B). We compared the co-accessibility of CREs within the same module versus across modules at the single-cell level by computing the Pearson correlation of DNA accessibility across individual cells in a peripheral blood mononuclear cell (PBMC) scATAC-seq dataset, and found higher single-cell co-accessibility among CREs within the same module (Figure 1C). This was also visually apparent when inspecting the DNA accessibility profiles of PBMC cell types, where CREs were grouped according to their co-accessibility rather than purely based on genomic proximity (Figure 1D). To test whether REMO modules capture shared regulatory networks, we linked CREs to putative target genes by correlation-based peak-gene linkage using a collection of 10x Multiome datasets from 16 different human tissues. We then used logistic regression to model the probability of two CREs belonging to the same module based on whether they regulate the same gene, controlling for genomic distance between the CREs. This revealed that CRE pairs regulating the same gene were significantly more likely to be within the same module than distance-matched CRE pairs linked to non-overlapping gene sets (odds ratio ranging from 1.87 to 2.21 across all chromosomes, p < 2 x 10^-16^). We also observed that single-CRE modules were 1.84 times less likely to be linked to a gene, indicating that these CREs may be less likely to participate in functional regulatory networks.

### REMO provides a universal feature set for scATAC-seq analysis

To assess the performance of REMO as a set of features for scATAC-seq analysis we quantified the accessibility of REMO modules in a human PBMC multiome dataset assaying mRNA abundance and DNA accessibility within the same cell. The distribution of the REMO count matrix appeared to be more similar to that of the RNA count matrix, with higher count values more frequently observed compared to the peak count matrix (Figure 2A). As expected, the mean and variance of REMO features were highly correlated, but we observed some modules that appeared to deviate from the expected mean-variance relationship, similar to the pattern seen for highly variable genes for scRNA-seq data (Figure 2B), whereas the peak matrix displayed very little deviation from the mean-variance equivalence. We selected the top 20,000 most highly variable REMO modules based on the residual variance from a mean-variance LOESS model fit. We performed dimension reduction on the matrix of variable REMO modules using principal component analysis (PCA), and on the peak matrix using latent semantic indexing (LSI) ^6^. UMAP visualization of the cells revealed a similar separation of the different cell types when processing the dataset using REMO or the peak matrix, with cells annotated according to their gene expression profiles (Figure 2C).

**Figure 2:**
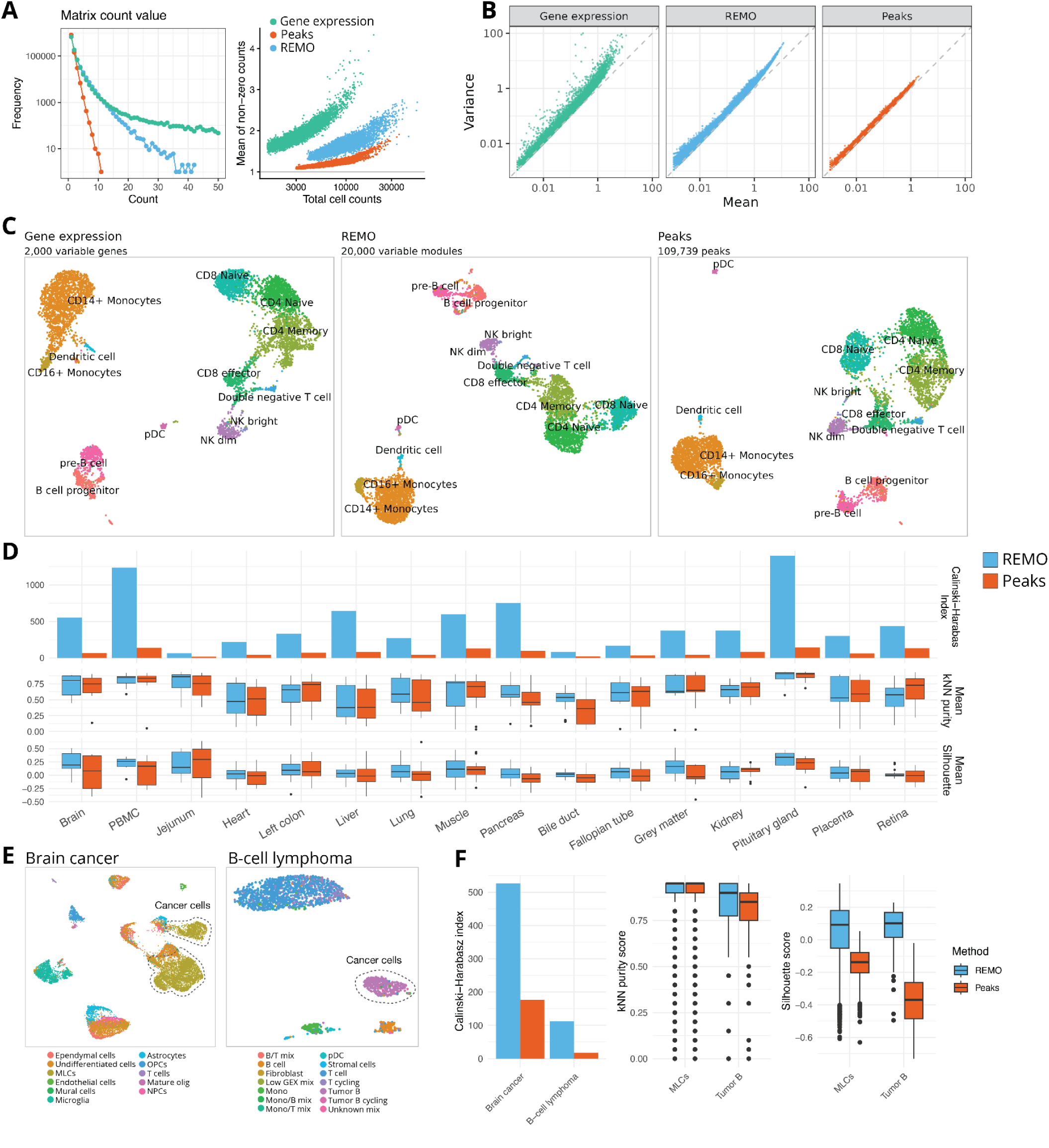
REMO provides a universal feature set for scATAC-seq data analysis. **(A)** Distribution of non-zero count matrix values for RNA, REMO, and peak count matrices in the PBMC multiome dataset. **(B)** Log-mean vs log-variance relationship for the RNA, REMO, and peak count matrices. **(C)** UMAP representation of PBMC multiome dataset cells processed using the gene expression assay (left), REMO (center), or peaks (right). In each plot, cells are colored according to their cell type label determined from the RNA assay. **(D)** Calinski-Harabasz index, mean kNN purity, and mean silhouette width for each cell cluster across 16 different multiome datasets, computed using the low-dimensional space from REMO or peak matrix processing. For each dataset cell clusters were determined by processing the RNA assay, independently from the DNA accessibility data. **(E)** REMO-based UMAP plots showing the 2-dimensional projection of cells from a brain cancer (pediatric posterior fossa ependymoma, MLCs: mesenchymal-like tumor cells) and B-cell lymphoma (small lymphocytic lymphoma of the lymph node: Tumor B) patient. Cells are annotated according to author-provided cell labels, with the cancer cells circled by a dashed line. **(F)** Calinski-Harabasz index, mean kNN purity score and mean silhouette for cancer cells in the brain cancer and B-cell lymphoma datasets, for REMO and peak-based data processing.

To quantitatively compare the performance of REMO and peak-based data processing across a more comprehensive collection of human tissues, we gathered published multiome data for 16 different tissues and repeated a similar analysis for each dataset using both REMO and dataset-specific peak calling (Supplementary Figure 1, Supplementary Table 1). We compared agreement between the REMO-based and peak-based low-dimensional embeddings and the RNA-derived cell cluster labels for each tissue using three different metrics: the Calinski-Harabasz index (CH index), the mean k-nearest neighbor (kNN) purity score, and the mean silhouette score (Figure 2D). The CH index measures the ratio of between-cluster separation to within-cluster dispersion (one value per dataset; higher is better). Similarly, the silhouette measures the proximity of each cell to other cells of the same cluster compared to those of the neighboring cluster (one value between -1 and +1 per cell, higher is better). The kNN purity score measures the fraction of each cell’s neighbors that belong to the same cluster as that cell (one value between 0 and 1 per cell, higher is better). The kNN purity score and mean silhouette were similar for peaks and REMO for most tissues, indicating that the overall cell neighborhoods were similar when using the two different sets of features, although we observed some tissues where higher purity was obtained using REMO. However we observed higher CH indices for all tissues when using REMO compared to the peak matrix. This indicates that clusters were more compact and better separated in the REMO PCA space compared to in the peak LSI space. Importantly, this analysis re-used the same REMO modules across all tissues, demonstrating that high performance can be obtained with REMO as a universal set of features for scATAC-seq analysis, whereas each peak matrix involved *de novo* peak calling on the individual dataset and a much larger total number of features.

As the REMO modules were identified using healthy human cell states, we also wanted to assess the application of these features to scATAC-seq data containing disease-associated cell states. We performed a similar analysis on datasets from a brain cancer patient and B-cell lymphoma patient. For each dataset, we focused on the separation of the cancer cells from other cells in the dataset in the low-dimension space. In both datasets we observed similar neighborhood composition of cells in the low-dimension space, but greater separation of the disease cells from other cells in the dataset (as measured by the mean silhouette score and CH index) when using REMO features (Figure 2E, F). This was consistent with results from the 16 healthy tissues, and indicates that REMO can be applied to both healthy and disease samples and achieves equivalent or greater performance than peak calling in projecting cells into a low-dimensional space.

### Highly scalable analysis of scATAC-seq data with fragtk and REMO

As REMO reduces the total number of features used, we reasoned that the reduced matrix size should enable a more computationally scalable analysis. However, REMO still requires that >1.5 million genomic regions are quantified per cell. This presents a challenging problem that some of the current scATAC-seq software tools handle poorly, and requires summing accessibility values across non-contiguous genomic regions for each cell, something that is not possible with current tools. To address this gap we developed a new software package for scATAC-seq data quantification, named fragtk, with an emphasis on speed and memory efficiency.

To assess the performance of fragtk we obtained a published scATAC-seq dataset of 600,000 cells from the whole adult human body ^7^. We downsampled this dataset to different numbers of cells and assessed the time and memory required to quantify a set of 343,707 peaks in each cell using four different software tools: Signac ^6^, SnapATAC2 ^8^, BPcells ^9^, and fragtk. Fragtk had a faster runtime than all other tools across the range of dataset sizes (Figure 3A, left panel). In some cases these time savings were substantial (2 hours faster quantification than SnapATAC2 on 600,000 cells). Reductions in the amount of memory required were also observed in comparison to SnapATAC2 and Signac, with both BPcells and fragtk requiring a small memory footprint that did not substantially increase with increasing numbers of cells (Figure 3B, right panel).

**Figure 3:**
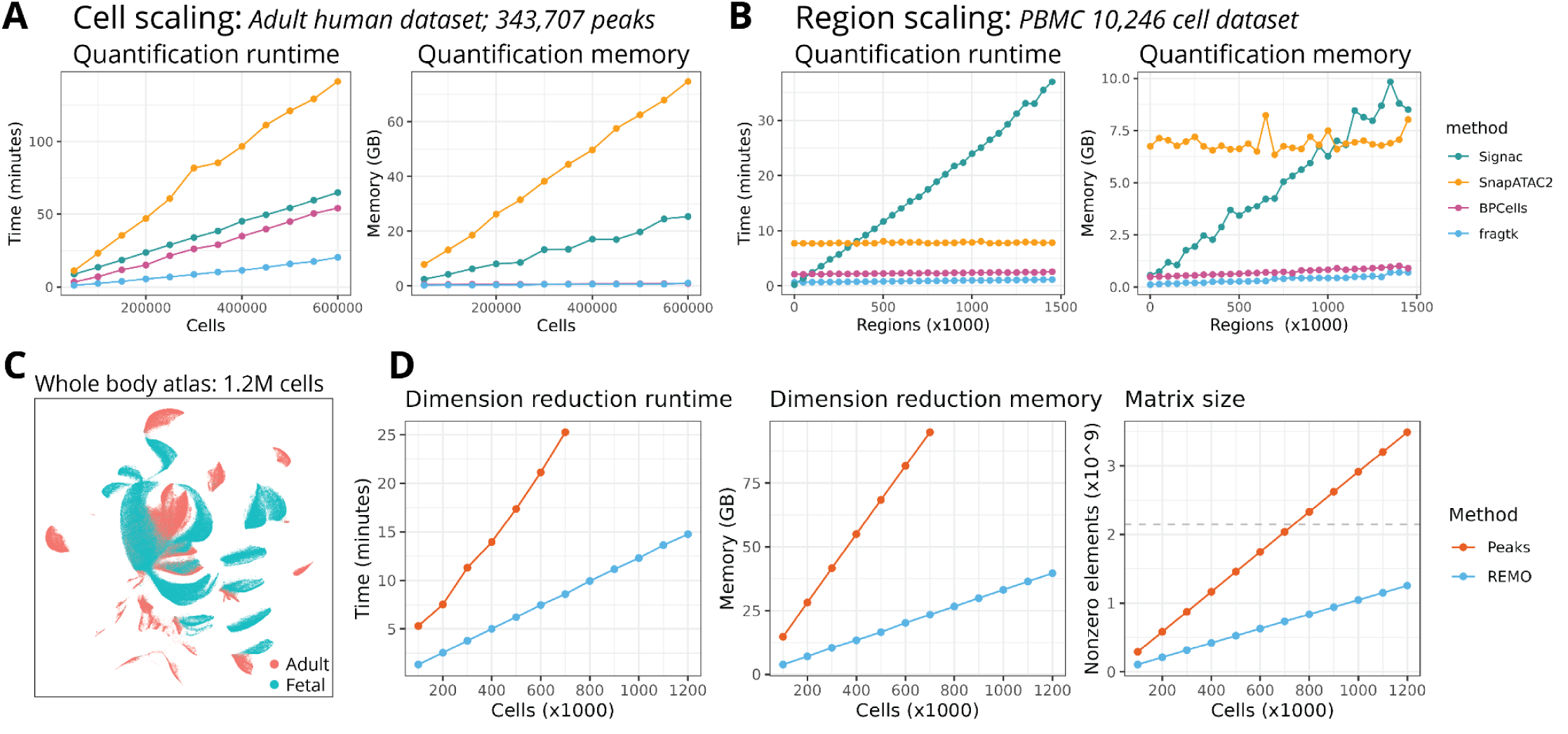
Scalable data quantification and dimension reduction. **(A)** Runtime and memory use for quantification of 343,707 peak regions across different numbers of cells sampled from the 600,000-cell human chromatin accessibility atlas dataset, for Signac, SnapATAC2, BPCells, and fragtk. **(B)** Runtime and memory use for quantification of different numbers of genomic regions for 10,246 cells, for Signac, SnapATAC2, BPCells, and fragtk. **(C)** UMAP representation of the combined fetal and adult human chromatin accessibility atlas complete dataset (1.2 million cells) processed using REMO. **(D)** Runtime, memory, and matrix size (number of nonzero entries in the sparse matrix) comparison for REMO and peak matrix processing of different random cell subsets ranging from 100,000 to the full 1.2 million cell atlas dataset. The dashed line represents the size limit of a 32-bit sparse matrix. Values not shown for peak matrix processing indicate that the matrix was unable to be loaded in R as it exceeded the maximum size of a 32-bit sparse matrix. For each set of downsampled cells, the time and maximum memory required to normalize the data and compute a low-dimensional space was recorded.

We next assessed the scalability of quantification methods in terms of the number of genomic regions quantified, using a smaller dataset of 10,246 PBMCs. We quantified different subsets of the collection of >1.5 million CREs, ranging from 1,000 to 1,500,000 genomic regions. Signac performed poorly in this analysis and displayed increasing runtime and memory requirements with increasing numbers of genomic regions. SnapATAC2, BPcells, and fragtk had runtimes and memory requirements that remained relatively constant with increasing numbers of regions, with fragtk having the fastest runtime and lowest memory footprint in all cases (Figure 3B).

To assess the scalability of scATAC-seq data processing using REMO in comparison to peaks, we combined the adult human scATAC-seq atlas dataset with a scATAC-seq atlas from fetal tissues ^10^, resulting in a combined dataset of 1.2 million cells (Figure 3C). We measured the runtime and memory requirements to perform data normalization and dimension reduction on different subsets of this dataset using the peak matrix (678,387 peaks) or the top 20,000 highly variable REMO modules. To enable memory-efficient feature selection, we also developed a new software tool (spars) to perform disk-based feature selection and subsetting of sparse matrices. With 700,000 cells (the highest number able to be processed using the peak matrix), we observed 3x faster runtime (8 minutes vs 25 minutes) and 4x lower memory use (23 GB vs 94 GB) with REMO compared to the peak matrix (Figure 3D). Using REMO we were able to quantify the full dataset of 1.2 million cells without reaching the maximum size limit of a 32-bit sparse matrix, and completed dimension reduction in 14 minutes. This was less than the time taken to process 500,000 cells using the peak matrix.

### Annotation of scATAC-seq data via cell ontology term enrichment

Another major challenge in the analysis of scATAC-seq data is the lack of interpretable information associated with the assay features. This is compounded by studies using different peak sets, preventing the accumulation of data associated with genomic features in the biomedical literature over time. In contrast to gene expression analysis, we lack information associating features measured in the assay with cell types. To address this limitation we annotated the top cell types accessible for each REMO module using a collection of 144 cell types curated from published scATAC-seq data from the whole body, harmonized to Cell Ontology (CL) terms ^11^. For each REMO module we identified a list of cell types in which CREs in the module were most highly accessible, weighted by their cell type specificity, and annotated the modules with these CL terms. To enable a subset of cell types to be selected according to a tissue of interest, we mapped cell types to a list of tissues in which those cell types are present by combining information from the Human Reference Atlas Cell Type Annotations ^12^ for Azimuth ^13^, CellTypist ^14^, popV ^15^, and manual assignment.

To evaluate the accuracy of cell annotation by CL term enrichment testing we analyzed multiome datasets from the brain, PBMCs, and pancreatic islets ^16–18^. These datasets were not used in the annotation of REMO modules with CL terms. For each dataset we annotated clusters using label transfer from an annotated scRNA-seq reference using the gene expression data ^13,19^. Next, we performed differential testing for each cell cluster using the REMO modules and applied ontology term enrichment testing to identify CL terms enriched in REMO modules specific to each cluster ^20^. This was done either using all CL terms (whole body mapping) or a subset of terms for the query tissue (tissue mapping) (Figure 4A). Overall we observed high agreement between cell type annotations derived from a term enrichment test with REMO modules and when using the gene expression information (Figure 4B, Supplementary Table 2). This shows how cell ontology term enrichment analysis can provide a simple approach to return biologically interpretable information for scATAC-seq datasets, analogous to lists of cell-type-specific marker genes in transcriptomic analysis, and can provide an accessible starting point for more refined cell type annotation.

**Figure 4:**
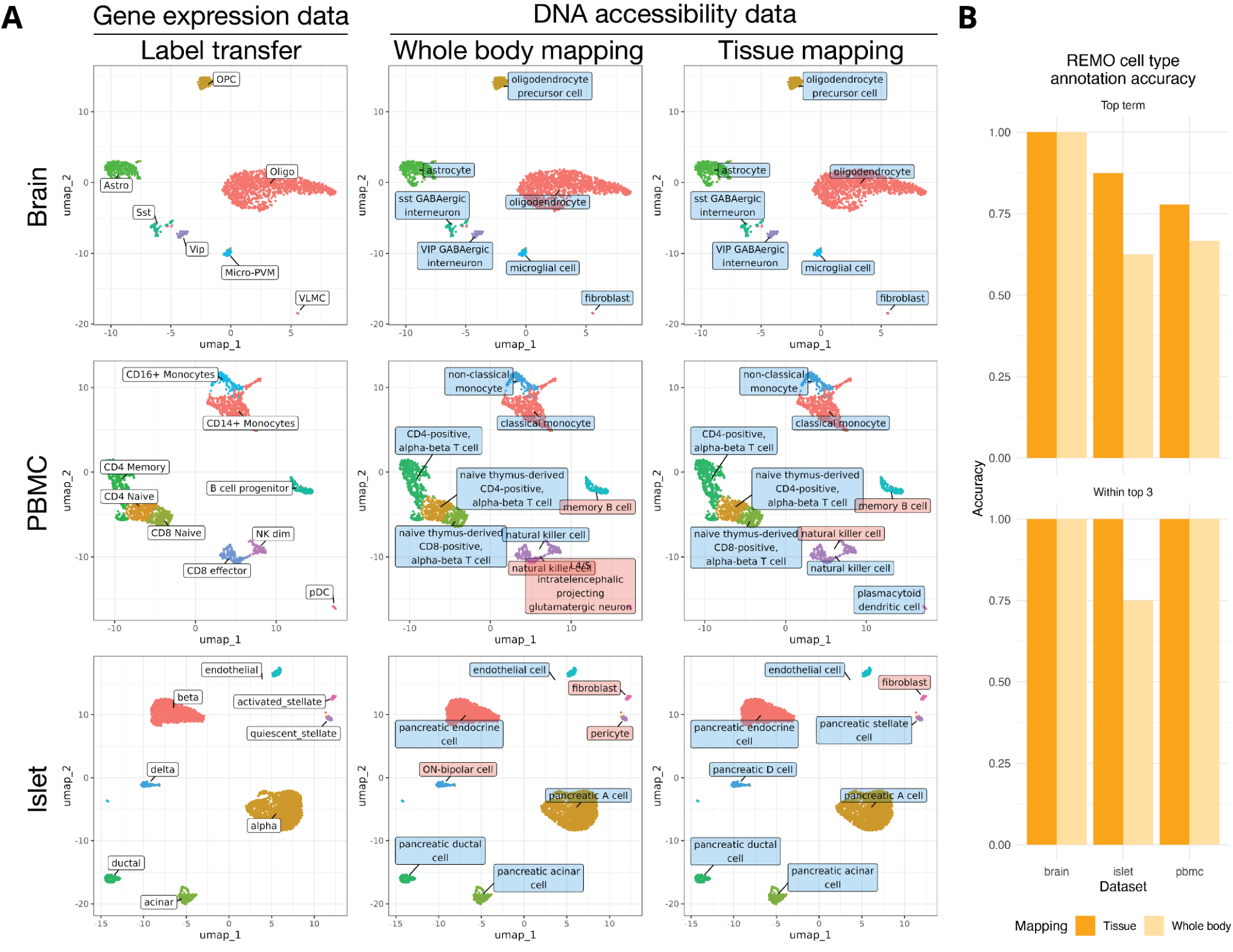
REMO cell ontologies enable annotation of scATAC-seq data. **(A)** Annotation of brain, PBMC, and islet cell clusters. Each dataset was projected into a two-dimensional visualization using PCA and UMAP on the REMO counts. Cell clusters were annotated by reference mapping using the gene expression data (first column) or by cell ontology term enrichment testing, either mapping to all cells (whole body mapping, second column), or cells in the specific tissue (brain, blood, or pancreas; third column). Correctly annotated cell clusters are shown with blue labels, incorrect annotations are shown with red labels. Cell annotations derived from the gene expression data are shown with white labels. **(B)** Accuracy of cell type annotation for each dataset when mapping to the whole body or to the tissue of interest. “Top term” indicates that the correct cell type was the top enriched term. “Within top 3” indicates that the correct cell type was within the top three enriched terms. Accuracy is the fraction of cell clusters matching the gene expression-derived cell type label.

## Discussion

The re-use of public scATAC-seq data is complicated by a lack of common features, making the sharing of count matrices of little practical value. To compare their data to other published datasets, researchers instead need to access published data in the form of a fragment file, BAM file, or FASTQ files, and perform additional data processing steps. This presents substantial barriers to data re-use, especially as raw data from human subjects often must be placed in controlled-access data repositories. Here, we sought to address this major barrier by developing a set of universal features for human scATAC-seq analysis. While we have demonstrated several major advantages to using REMO (including improved separation of cell states in a low dimensional space, speed and memory efficiency, and data interpretation) we expect that the greatest benefit of broad adoption of REMO will the the accumulation over time of publicly available scATAC-seq count matrices quantifying a common set of features. This will be an incredibly valuable resource for the field as it will allow individual studies to place their findings in the context of other datasets. Furthermore, it will enable building an atlas-scale data repository for single-cell chromatin accessibility data that aggregates information across many individual studies. Such a repository will be a valuable resource with many potential applications, and equivalents already exist for single-cell transcriptomic data ^21^.

Alongside the development of REMO we also report the development of new quantification software for scATAC-seq data, fragtk. Fragtk is fast and memory-efficient, outperforming existing tools in these respects, and provides an option to perform paired insertion counting ^22^. In developing fragtk we also aimed to create a tool that is user-friendly and simple to install. As a result, installation is as simple as downloading a single pre-compiled binary file, and no dependencies are needed. Open-source Rust code is also made available. To facilitate the analysis of scATAC-seq data using REMO we have also incorporated new functionality into our open-source R package Signac ^6^, and provide a new R package containing all of the REMO module information including cell ontology terms (see Code Availability below).

Here we provide an initial REMO set for the human hg38 genome. We see this as a starting point, and envision that future updates to the catalogue of CREs in the human genome, and other data sources, will enable new and updated REMO versions to be developed in the future with improved performance when applied to single-cell data. We also envision that similar efforts can be undertaken for other genomes, particularly the mouse and other key model species, to identify REMO modules for model species.

## Methods

### Grouping regulatory element modules

A comprehensive set of CREs for the human genome was constructed by combining the candidate cis-regulatory elements (CREs) from ENCODE SCREEN v3 ^3^ with a set of previously-reported consensus ATAC-seq peaks (cPeaks; ^4^). Regions from the ENCODE CREs and cPeaks were combined using GenomicRanges in R ^23^. For overlapping regions between cPeaks and ENCODE, the cPeaks coordinates were used. This produced a final set of 1,520,441 genomic regions.

To provide an initial grouping of CREs based on their biochemical activity across a broad range of tissue types, we quantified signal in each CRE across a set of 4,157 histone and transcription factor ChIP-seq datasets available from ENCODE (Supplementary Table 3) ^3^ using the bigwigaverageoverbed function from the bigtools package ^24^. This matrix was split by chromosome and CREs for each chromosome processed separately. Outlier values greater than the 90th percentile for each dataset were clipped to the 90th percentile value. The CRE-by-experiment data matrix was normalized with edgeR using the trimmed mean of M-values (TMM) as the normalization factor ^25^. CREs were projected into a low-dimension space by performing PCA on the normalized matrix with the irlba R package ^26^. A k-nearest neighbor (kNN) graph was then constructed by finding the first 50 neighbors for each CRE in the 50-dimension PCA space ^27^, and converted into a shared nearest neighbor (SNN) graph by computing the neighborhood overlaps (Jaccard similarity between CREs and each nearest neighbor). Clusters were identified using the Leiden clustering algorithm in the igraph R package with the modularity objective function ^28,29^.

Within each broad-level cluster of CREs we performed subclustering using DNA co-accessibility of CREs, the DNA physical contact probability, and the genomic distance between CREs. We computed a co-accessibility-by-contact score *S* between each pair of CREs *i* and *j*:

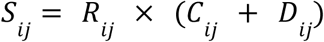

Where *R*_*ij*_ is the Pearson correlation of DNA accessibility between CRE *i* and CRE *j*, with negative values set to zero, computed using a large collection of pseudobulk cell clusters from scATAC-seq datasets (see below); *C*_*ij*_ is the normalized HiC contact score between the pair of CREs, with HiC contacts for the chromosome min-max scaled between 0 and 1 at 5 kb resolution; *D*_*ij*_ is the genomic distance transformed using a Gaussian kernel:

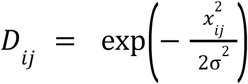

Where *x* is the genomic distance between the pair of CREs in base pairs, and σ is the Gaussian kernel bandwidth (set to 500,000). After computing *S* between each CRE pair, a weighted graph was constructed by taking the top 30 edges for each node (CRE) ranked by their score *S*, with edge weight set to *S*. We then identified clusters in the weighted graph using the Leiden community detection algorithm with the Constant Potts Model objective function (cluster_leiden function in igraph; resolution=0.5, objective=’CPM’) ^28,29^. This was repeated for each chromosome to define the complete set of REMO modules. The Constant Potts Model was chosen as it tends to perform better than modularity at identifying small, densely-connected communities.

### Chromatin accessibility datasets for co-accessibility estimation

To calculate the co-accessibility between CREs we constructed an extensive CRE accessibility dataset spanning many human tissues by compiling published scATAC-seq and multiome datasets. This consisted of a large-scale chromatin accessibility atlas datasets, as well as an aggregation of published datasets for individual tissues (Supplementary Table 4). All datasets were mapped to the hg38 reference genome.

#### Atlas datasets

We obtained data from large-scale chromatin accessibility atlas studies. This included a whole body single cell dataset with 222 cell clusters ^7^, a developmental brain atlas with 135 cell clusters from the CATlas website ^30^, and an adult human brain dataset with 107 cell clusters from the Neuroscience Multi-omic Archive (NeMO) website ^31^. For each atlas dataset bigwig files of all available cell clusters were downloaded and values overlapping CREs were quantified using the bigwigaverageoverbed function from bigtools ^24^.

#### Individual human tissue datasets

A pseudobulk chromatin accessibility matrix was created for cell clusters in a collection of human tissues (adrenal, muscle, esophagus, fetal heart, heart right ventricle, left colon, liver, blood) using scATAC-seq and 10x Multiome datasets from 10x Genomics and ENCODE ^3^. We processed these datasets and generated a pseudobulk CRE accessibility matrix using separate pipelines for immune datasets and other human tissues. The immune pseudobulk matrix represented accessibility information of 98,584 cells from 16 PBMC cell types. The human tissue pseudobulk matrix represented accessibility information of 106,027 cells from 72 cell clusters.

#### Immune datasets

The pseudobulk immune atlas was created by combining 16 publicly available scATAC-seq and 10x Multiome datasets from 10x Genomics. For each sample the fragment file, peak coordinate BED file, and filtered feature matrix (for multiome datasets only) were obtained from the 10x Genomics website. For multiome datasets, the RNA assay was depth normalized and log transformed (NormalizeData function in Seurat), and dimension reduction performed using feature selection and PCA (FindVariableFeatures, ScaleData and RunPCA functions). Cell types were annotated for multiome datasets by transferring cell type labels from an existing PBMC reference dataset ^13^ using the Seurat label transfer methods (FindTransferAnchors followed by TransferData functions ^19^). Single-cell ATAC and multiome datasets were integrated using reciprocal LSI projection (FindIntegrationAnchors function with reduction=”rlsi”, dims=2:30, followed by IntegrateEmbeddings function). After data integration, scATAC-seq cells were annotated via label transfer with the previously annotated multome cells as a reference object, and using the DNA accessibility information for anchor-finding (reduction.method=”lsi” in FindTransferAnchors followed by TransferData function).

#### Non-immune datasets

A human pseudobulk DNA accessibility atlas was created using publicly available scATAC-seq datasets across seven human tissues (adrenal, psoas muscle, esophagus, fetal heart, adult heart right ventricle, left colon, liver), with a total of 19 samples with a minimum of 2 samples per tissue. Fragment files for each sample were obtained from the ENCODE consortium database ^3^. Peak calling was performed using the callpeak function from MACS2 ^32^ with the following parameters: --nomodel --shift -100 --extsize 200. For each individual tissue, a common peak set was constructed by collapsing overlapping peak ranges using the “reduce” function from GenomicRanges R package ^23^. Remaining peaks shorter than 20 bp or longer than 10,000 bp, or that overlapped with genomic blacklist sites ^33^ were excluded. A peak-by-cell count matrix was constructed for each dataset using the fragtk matrix command.

A Seurat object was created for each sample using the peak count matrix. Quality control metrics such as the number of ATAC fragments mapped to peaks, the number of RNA UMIs detected (multiome datasets only), percentage of peak region fragments, TSS enrichment score, and nucleosome signal were computed and low-quality cells based on these QC metrics were excluded (Supplementary Table 5). Dimension reduction for the DNA accessibility assay in each dataset was performed using LSI (RunTFIDF and RunSVD functions in the Signac package ^6^). To integrate multiple scATAC datasets of the same tissue we used Seurat integration with reciprocal LSI projection (FindIntegrationAnchors function with reduction=”rlsi”, dims=2:30) ^19^. Low dimensional cell embeddings across datasets were integrated using the IntegrateEmbeddings function with parameters dims.to.integrate=1:30. Next, a k-nearest neighbors graph was constructed from the integrated LSI embeddings using dimensions 2 to 30 (FindNeighbors function with parameters nn.method=”annoy”, k.param=20, dims=2:30). Clusters were detected using the smart local moving (SLM) algorithm ^34^.

#### Pseudobulk DNA accessibility matrix construction

For the integrated immune dataset, we retained cell clusters with greater than 500 total cells for construction of a pseudobulk DNA accessibility matrix. The scATAC-seq fragment files for each dataset were split into separate files for each cell cluster using the SplitFragments function from Signac. A bedGraph file of per-base coverage was generated using the genomecov function from bedtools ^35^, and converted to a BigWig file using the bedgraphtobigwig function from bigtools ^24^. A pseudobulk DNA accessibility matrix was then constructed by computing the accessibility for each CRE region for each pseudobulk cell cluster using the bigwigaverageoverbed function from bigtools, and individual accessibility vectors combined into a matrix. For datasets from non-immune tissues, a pseudobulk accessibility matrix was constructed by first constructing a CRE-by-cell accessibility matrix in R using the FeatureMatrix function in Signac. This matrix was collapsed to a CRE-by-cluster matrix by performing matrix multiplication with a binary cluster-by-cell membership matrix created using the BinaryIdentMatrix function from Signac, and normalized by the number of cells per cluster.

### HiC datasets

A collection of 45 deeply sequenced Hi-C datasets were obtained from ENCODE (Supplementary Table 6). Hi-C contact matrices were loaded into R using the “straw” function from the strawr R package ^36^. The observed over expected (oe) values with “SCALE” normalization were used to estimate contact probabilities between genomic loci, with a bin size of 5,000 base pairs. To account for outliers, Hi-C values above the 90th quantile were clipped to the 90th quantile value. Hi-C contact values were averaged across the 45 datasets and scaled from 0 to 1 (min-max scaling) to produce a final contact probability for use in calculating co-accessibility-by-contact scores. Hi-C contact estimation was limited to CRE pairs within 1 million base pairs of each other.

### REMO performance analysis

To evaluate the performance of DNA accessibility assay data processing using REMO or using peaks, we utilized a collection of single-cell multiome data to compare the agreement between RNA-based and chromatin accessibility-based cellular analysis. We used publicly available multiome datasets for 16 healthy human tissues (brain, PBMCs, jejunum, fetal heart, left colon, liver, lung, muscle, pancreas, bile duct, fallopian tube, brain grey matter, kidney cortex, pituitary gland, placenta, retina) and two disease datasets (B-cell lymphoma and brain cancer). Each dataset was processed using the same processing steps and the performance of REMO and peaks compared using the gene expression-derived cell clustering labels. The sample information for each dataset can be found in Supplementary Table 1. Three Seurat objects were created for each tissue multiome dataset: an RNA object (gene expression), a peak object (chromatin accessibility using peak counts), and a REMO object (chromatin accessibility using REMO counts). Initial raw data processing for each 10x Multiome dataset was performed using the 10x Genomics Cell Ranger ARC software version 2.0.2, scATAC-seq data was processed using Cell Ranger ATAC version 2.1.0. Reference genome version 2020-A (GRCh38, equivalent to GENCODE v32/Ensembl98) was used for all datasets.

#### ENCODE datasets

For datasets originating from ENCODE (bile duct ENCSR871JTA, fallopian tube ENCSR420IUS, heart ENCSR302EOG, left colon ENCSR925IHI, liver ENCSR728OVE, lung ENCSR264JIX, muscle ENCSR851GBP, pancreas ENCSR233SQG, placenta ENCSR694BTU), we obtained FASTQ files from the ENCODE website and processed them using the cellranger-arc count function to generate fragment files and filtered feature matrices, mapping to the GRCh38 reference genome.

#### GEO datasets

##### Healthy datasets

For datasets available from published studies with data available on NCBI GEO (grey matter sample “PD003” GSE193240 ^37^; kidney cortex sample “AJDV174” GSE232222 ^38^; pituitary gland sample “Sample 1P-M” GSE178453 ^39^; retina sample “Multi_Fetal_23w1d_FR” GSE268630 ^40^), we used the fragment files and filtered feature matrix provided by the authors, as they were already processed using the 10x Genomics cellranger-arc software and mapped to the GRCh38 genome.

##### Disease datasets

For the brain cancer dataset (pediatric ependymoma, samples “I2”, “I3”, “I4”, “I5”, “M7”, and “M8”, <u>GSE206579</u> ^41^), fragment files for each sample was generated using cellranger-atac. A Seurat object containing 6 scATAC-seq samples with cell type annotations was provided by the original authors ^41^. We extracted the cell barcodes, peak coordinates, and cell type labels for further analysis from the published Seurat object. No additional filtering of cells was performed. To create the peak count matrix we used the fragtk matrix command with the author-provided peak coordinates as the input BED file. For the B-cell lymphoma dataset we downloaded the fragment file, peak count matrix, and loupe object containing cell type annotations from 10x Genomics website ^42^.

#### Count matrix generation

The fragtk count command was used to count the total number of fragments per cell. For each dataset, the top 10,000 cells with the highest number of fragment counts were selected for initial quantification and subject to further quality control filtering. Peak calling of the ATAC fragments was done using MACS2 ^32^ callpeak function with the following parameters: --nomodel --shift -100 --extsize 200. The number of peaks identified per tissue ranged from 43,378 to 229,286. To create the counts matrix for the peaks and REMO features, the fragtk matrix command was used with either the peaks or REMO modules as the input BED file, with the --pic option set to perform paired insertion counting ^22^, and the --group option set for the REMO features to perform grouped quantification. For the RNA assay, the filtered gene expression matrix from cellranger-arc was used.

#### Quality control

Multiome datasets were processed using Seurat (v5.3.0) and Signac (v1.16.9003) in R. Quality control (QC) metrics such as the number of ATAC fragments mapped to peaks, the number of RNA UMIs detected, TSS enrichment score, and mitochondrial RNA content were calculated and cells were excluded based on sample-specific QC thresholds (Supplementary Table 1). Doublets were identified using the “scDblFinder” function from scDblFinder (v1.16.0) with default parameters ^43^, cells that were not classified as the “singlet” class were removed from the dataset.

#### RNA object processing

For the RNA objects, the gene expression count matrix was log normalized with a scale factor of 10,000 (NormalizeData function in Seurat). Variable genes were selected using the FindVariableFeatures function in Seurat (selection.method=”vst”). Dimension reduction was performed using principal component analysis (PCA) with the ScaleData and RunPCA functions in Seurat. Cell clustering was performed for each dataset by constructing a neighbor graph and performing graph-based clustering (FindNeighbors with reduction=”pca” followed by FindClusters with algorithm=3, resolution=0.5).

#### REMO object processing

For REMO objects, the REMO count matrix was log normalized as described for the RNA assay. Variable REMO modules were selected using the FitMeanVar function in Signac v1.16.9003, selecting the top 20,000 features for each dataset. This function fits a local regression curve (LOESS) to the log-mean and log-variance of each feature, using a set of features evenly downsampled across the range of mean and variance values. Variable features are selected based on the highest residual variance from the regression model. This is very similar to the FindVariableFeatures function in Seurat, but is more scalable to larger numbers of features through the use of downsampling for the model fit. Dimension reduction was performed using PCA using the RunSVD function in Signac with “pca=TRUE”. This setting allows for computation of PCA components using the Rspectra package without storing the scaled and centered data matrix, reducing the memory required ^44^. A kNN graph was constructed using the FindNeighbors function in Seurat with reduction=”pca”.

#### Peak object processing

For peak matrix data processing, features were selected using the FindTopFeatures function in Signac with default parameters. Dimension reduction was performed using latent semantic indexing (LSI; RunTFIDF function followed by RunSVD function in Signac). A kNN graph was constructed using the FindNeighbors function in Seurat with reduction=”lsi”.

#### Count distribution comparison

To visualize the count distribution for REMO counts, peak counts, and gene expression counts for the PBMC multiome dataset (above) we sampled 1,000,000 non-zero elements from each count matrix at random and plotted the frequency of each integer value for each count matrix. We also plotted the relationship between the mean of non-zero counts and the total counts detected for the cell, as suggested by Kwok et al. ^45^.

#### CRE-gene linkage analysis

To assess whether CREs regulating the same gene were more likely to be placed within the same REMO module, we linked each CRE to a potential set of regulated genes using the 16 multiome datasets (above). The LinkPeaks function in Signac ^6,46^ was modified such that instead of excluding only the peaks within a certain distance from the longest TSS, the TSS of all transcripts for a given gene within the gene.coords argument were considered. This was run across each dataset with the following parameters: distance = 5e+05, min.distance = 500, min.cells = 10, sample = 200, pvalue_cutoff = 1, score_cutoff = 0. The GENCODE v32 basic annotation file was supplied as gene.coords, as this was the gene annotation used for gene expression quantification in the multiome datasets. Results were combined across tissues by taking the highest Pearson correlation value for each significant (p < 0.05) CRE-gene link across all tissues.

To determine the likelihood that CREs linked to the same gene colocalise within the same module, a statistical modelling approach was used to control for distance. For this analysis, CREs not linked to any gene were excluded, and only pairs of CREs within 1 Mb of each other (the maximum possible span between two candidate CREs linked to the same TSS by LinkPeaks) were considered. For each chromosome, we then fitted a logistic regression model via the glm function in R to predict the probability of each CRE pair *i* and *j* residing in the same module given their gene linkage. This was parameterised using the log distance between the CRE pair *log*(*d*_*ij*_), and a binary variable *X*_*samegene*_ for whether their linked gene sets shared at least one element. The beta coefficient β_2_ of the model and its associated p-value were noted.

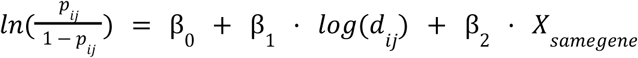

To assess whether single-CRE modules were more or less likely to be linked to genes, a simple Bayesian approach was applied. From a frequency table of the number of CREs that were linked *N*(*C_linked_*), or unlinked *N*(*C_unlinked_*) and located in a single-CRE module *N*(*M_single_*) or multi-CRE module *N*(*M_multi_*), we computed the probability that a CRE was unlinked given membership in a single-CRE module. This was compared to the base probability that a CRE was unlinked.

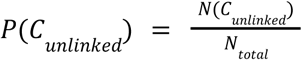

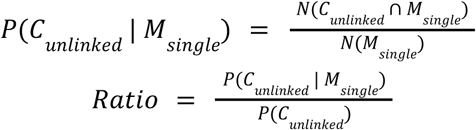

#### Dimension reduction performance evaluation

To compare the performance of REMO and peaks in projecting cells into a low-dimension space, we focused on measuring the level of agreement with gene expression-derived cell cluster labels. For each tissue, we identified cell clusters using the RNA object processing described above. We independently processed the DNA accessibility assay using REMO or peak matrix quantification and obtained low-dimensional cell embeddings, and computed performance metrics using the RNA-derived cluster labels and the assay-specific low-dimensional space. We computed the Silhouette score using the silhouette function from the cluster v2.1.8.1 R package ^47^. To account for unbalanced cell type proportions, we computed the mean Silhouette per cell cluster. We computed the Calinski-Harabasz (CH) index using the calinhara function from the fpc R package ^48^. We computed a k-nearest neighbor (kNN) purity score as the fraction of a cell’s nearest neighbors belonging to the same cell cluster classification as the query cell ^6^. This was computed by identifying the 20 nearest neighbors for each cell using the nn2 function from the RANN R package ^49^. Again, to account for unbalanced cell type proportions we computed the mean kNN purity per cell cluster.

#### Disease dataset analysis

To assess the ability of REMO to separate disease-associated cell states that were likely not present in the collection of data used to identify REMO modules, we obtained publicly available datasets for B-cell lymphoma (10x Multiome data) and pediatric posterior fossa ependymoma (scATAC-seq data), each with published cell type labels ^41,42^. We assessed the separation of disease-associated cells from other cells in the dataset in the low-dimensional space. We computed the silhouette score and kNN purity score as described above, but focused our analysis on the disease cell states rather than all cell clusters in the dataset. The CH index was computed for the whole dataset, as this provides one metric per dataset rather than per cell.

#### Scalability analysis

We combined scATAC-seq datasets from the adult ^7^ and fetal ^10^ human tissues to create a dataset of 1.2M cells. This was used to assess the scalability of data processing using peak counts and REMO counts. As data from the fetal tissues were originally mapped to hg19, we first downloaded the author-provided fragment files and lifted these over to the hg38 assembly using liftover and the hg19tohg38 chain file ^50^. Fragment files for each tissue type were combined into a single fragment file each for the adult and fetal datasets, appending the tissue of origin to the cell barcode, and the combined file position-sorted and bgzip-compressed using the unix sort command and htslib (v1.20). We extracted peak regions from the author-supplied Seurat object and lifted them over to hg38 using the hg19tohg38 chain file and the liftOver function in the rtracklayer R package, keeping uniquely lifted-over coordinates with length >200 bp, and removing any regions that intersected the set of unified genomic blacklist sites for hg38 ^33^. For the adult human tissues, we combined fragment files from individual tissues into a single file using the approach described above. Peak regions were identified using the combined fragment file by running MACS3 with the following parameters: -f FRAG –nomodel --nolambda -g 2.9e9 --extsize 200 --shift -100 ^32^. Peak regions for fetal and adult datasets were combined using the reduce function in GenomicRanges, filtering the resulting regions to include those on standard chromosomes, greater than 300 bp in length, less than 5,000 bp in length, and not overlapping any genomic blacklist regions for hg38 ^23,33^. This resulted in a set of 678,387 peak regions. Code to reproduce these steps is available at: https://github.com/stuart-lab/human-chromatin-atlas.

We downsampled cells from each dataset (adult and fetal) from 50,000 to 600,000 cells by sampling cell barcodes at random from the complete set for each dataset. A full count matrix for all cells was quantified for peak regions and for REMO features using the fragtk matrix command, with the --pic option set to use paired insertion counting, and the --group option set for REMO feature to perform grouped quantification. The full count matrix was subset on-disk to each set of downsampled cells using the spars subset command. The subset matrices for fetal and adult datasets were loaded into R using the Read10X function in Seurat and concatenated using cbind in R to produce a dataset with equal cell numbers from fetal and adult at each downsampling level. A low-dimensional LSI projection was computed for peak counts using the RunTFIDF and RunSVD functions in Signac. A low-dimensional PCA projection was computed for REMO counts using the NormalizeData function in Seurat followed by the RunSVD function in Signac with pca=TRUE. Runtime and maximum memory was recorded using the benchmarking feature in snakemake ^51^.

### Development of fragtk and spars

#### Fragtk

We developed a fragment file toolkit (fragtk) to perform common operations on fragment files, the common data structure used for single-cell chromatin analysis. This toolkit was implemented using Rust and has four functions (as of v1.6): count, matrix, filter, and qc. The count function counts the total fragments per cell barcode contained within the fragment file, and has options to output a selected set of cell barcodes either by a minimum count threshold or by choosing the top *n* barcodes. The matrix function enables computation of a cell-by-feature matrix, given a list of cell barcodes to include and a set of genomic regions. Two counting strategies are enabled: insertion counting or paired insertion counting ^22^. We used the rust_lapper data structure, which uses a binary search on start positions, to efficiently store and query genomic ranges ^52^. The flate2 crate was used for decompression of the bgzip-compressed fragment file, and separate threads used for decompression and data parsing. The filter function enables subsetting of a fragment file according to a list of cell barcodes to include, simply by iterating through lines in the fragment file and discarding those not in the inclusion list. The qc function enables calculation of per-cell quality control metrics, including the TSS enrichment score ^3^, nucleosome signal score ^6^, total fragments on the mitochondrial genome, and total fragments for each cell.

#### Spars

We developed a Rust toolkit, spars, to perform common operations on sparse matrices stored in the matrix market format (the mtx file format commonly used in single-cell analysis), using a small memory footprint. The spars package has two functions (as of v1.3): stats and subset. The stats function enables computation of statistics for each row and column of the matrix (nonzero count, sum, mean, variance, standard deviation, min, max). The subset function allows rows and columns of the matrix to be subsetted from the full matrix. Each operation is performed by iterating over the mtx file lines (each line corresponding to a single non-zero element of the matrix), and so requires a very small amount of memory.

### Data quantification benchmarking analysis

To evaluate the runtime and memory requirements for scATAC-seq data quantification using different tools, we used the adult human chromatin accessibility atlas dataset described above ^7^. We downsampled the number of cells in the dataset from 50,000 to 600,000 (step size of 50,000) and created a fragment file for each downsampled set of cells using the fragtk filter command to simulate datasets of various sizes. For each downsampled dataset we quantified counts in the set of 343,707 peak regions using fragtk v1.6 (fragtk matrix function with --pic), Signac v1.16.0 (FeatureMatrix function) ^6^, SnapATAC2 v2.8.0 (import_fragments followed by make_peak_matrix function) ^8^, and BPCells v0.3.0 (peak_matrix function) ^9^. Runtime and maximum memory was recorded using the benchmarking feature in snakemake ^51^.

To assess how runtime and memory changed according to the number of genomic regions quantified, we used a smaller dataset of 10,246 human PBMCs from 10x Genomics ^53^. The dataset processed by Cell Ranger ATAC 2.1.0 was downloaded from the 10x Genomics website, and the list of included cells used in downstream quantification. We randomly sampled the set of 1,520,441 CREs (described above) from 1,000 to 1,500,000 (step size 50,000) and quantified a count matrix using each of the four software tools as described above.

### Module enrichment testing

#### REMO cell ontology annotation

We assembled a collection of pseudobulk cell type DNA accessibility data from a manually curated set of publicly available scATAC-seq data, including a whole body atlas dataset ^7^ and adult brain atlas ^31^, as well as collections of 10x Multiome data from PBMC (Supplementary Table 4, 5; sample_type=”immune”), heart, pancreas, lung, retina, and kidney (Supplementary Table 7). For the whole body atlas and brain atlas we used the author-provided cell type annotations and per-cell-type bigwig DNA accessibility files to compute a matrix of pseudobulk DNA accessibility within each CRE. For multiome datasets, we performed annotation of cell types using a combination of label transfer from an annotated reference and inspection of marker gene expression, and retained cell types with more than 100 cells. For all pseudobulk datasets we removed any cell types with zero values for >70% of CREs.

Cell type names were matched to a set of Cell Ontology (CL) terms ^11^. First, we identified semantically similar cell type names in the CL using Claude Opus 4.1, providing an initial automated mapping of the original cell type name to a candidate CL term, as well as a CL ID. The EBI OLS API service was used to look up the Claude-derived CL ID and verify that the ID was valid and corresponded to the reported CL term. All cell type names and CL terms were then manually checked and updated to a correct CL term when necessary. This resulted in a set of 144 unique CL terms.

We associated each CL term with a list of human tissues in which that cell type is found by compiling information from the Human Reference Atlas Cell Type Annotations ^12^. We downloaded the list of CL terms and the associated tissue for the Azimuth ^13^, CellTypist ^14^, and popV ^15^ reference datasets from the Human Reference Atlas website. CL terms in our list were matched to the reference datasets to create a list of tissues for each cell type. For CL terms that were not listed in the reference datasets, we manually assigned these terms to tissue types based on a literature search for each cell type. These results were stored in a JSON file of tissue names and the list of CL terms for the tissue, available in the REMO.v1.GRCh38 R package (see Data Availability below).

To annotate each REMO module with a set of CL terms, we first depth-normalized the pseudobulk accessibility matrix counts using TMM scaling factors ^54^. Next, we computed the tau specificity index τ for each CRE ^55^:

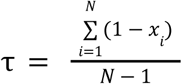

Where *N* is the number of cell types and *x*_*i*_ is the binarized accessibility of the CRE in cell type *i*. Accessibilities were binarized by identifying the 90% cumulative sum value for each cell type. This value was chosen as a binarization threshold, and CREs below this value set to zero and above the value set to one for the purpose of tau specificity index calculation only.

Depth-normalized pseudobulk accessibilities were transformed into decile values for each cell type (ranging from 0 to 1). For each REMO module we performed a weighted sum of the decile-transformed normalized accessibility of each CRE in the module, weighted by the CRE tau specificity index. As some cell types were present multiple times in the pseudobulk accessibility dataset (present in multiple tissues), the scores were averaged across multiple instances of the same cell type. The weighted sum values for each cell type were ranked for each REMO module, and the top 5 cell type terms recorded provided their weighted sum score was in the top 90% of scores across all cell type enrichment scores.

#### Cell type term enrichment testing

To compare cell annotation using label transfer with gene expression information versus CL term enrichment using REMO, we examined datasets from three different tissue types. We processed data from PBMCs ^18^, pancreatic islet (PANC-DB sample HPAP-175; ^16^), and brain ^17^ following the steps described in the previous section. Cell types were annotated using Seurat label transfer with using the following reference datasets: pbmc_10k_v3 ^19^, Azimuth v0.5.0 Human-Pancreas annotation ‘annotation.l1’ (pancreasref.SeuratData_1.0.0) ^13,56–61^, Azimuth v0.5.0 Human-Motor Cortex annotation ‘subclass’ (humancortexref.SeuratData_1.0.0) ^13,62^ for PBMC, pancreatic islet, and brain respectively. We used the FindNeighbors and FindClusters function from Seurat with resolution=2 to cluster each dataset, then annotated each cluster with the most frequent label transfer cell type with a prediction score of more than 0.5.

We used the cell type annotations from label transfer associated with each REMO cluster as a ground truth for cell annotation. For each cell cluster, the Seurat FindMarkers function was used to get differentially accessible REMO modules. A signed significance score *r* was calculated for each module *i* as:

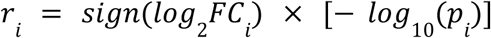

Modules were ranked according to *r*. Only modules included in the dataset’s variable features were included in the ranking. The fgsea function was then used to calculate set enrichment values using the module ranks and CL terms associated with each module ^20^. These results were sorted first by NES (decreasing), then by adjusted p-value (increasing) and any terms with adjusted p-value greater than 0.05 removed. These steps were performed using the EnrichedTerms function in Signac.

We mapped the top enriched term to each cell cluster and compared between the label transfer cell type names and CL terms from our REMO annotations (Supplementary Table 8). We measured accuracy as the fraction of cell clusters with matching label transfer and CL terms. Accuracy was calculated for the top-1 and within top-3 enriched terms, using all CL terms (whole body mapping) or a subset of terms for the query tissue (tissue mapping).

## Code availability

The fragtk and spars software packages are available from GitHub and crates.io: github.com/stuart-lab/fragtk, crates.io/crates/fragtk; github.com/stuart-lab/spars, crates.io/crates/spars.

Benchmarking analysis code for fragtk is available on GitHub: https://github.com/stuart-lab/fragtk-benchmark.

Code to reproduce regulatory element module identification is available on GitHub: github.com/stuart-lab/REMO.

Code to reproduce analyses shown in this manuscript is available on GitHub: github.com/stuart-lab/remo-manuscript.

## Data availability

REMO modules are available as a bgzip-compressed BED file through GitHub: https://github.com/stuart-lab/REMO.v1.GRCh38/blob/main/inst/extdata/REMOv1_GRCh38.bed.gz

REMO modules and metadata, including cell ontology term annotations, are also available as an R package: https://github.com/stuart-lab/REMO.v1.GRCh38

## Acknowledgements

This work is supported by the National Research Foundation, Singapore, under its NRF Fellowship program (T.S.; NRFF16-2024-0030). This work was supported by the A*STAR Computational Resource Centre through the use of its high performance computing facilities. This manuscript used data acquired from the database (https://hpap.pmacs.upenn.edu/) of the Human Pancreas Analysis Program (HPAP; RRID:SCR_016202; PMID: 31127054; PMID: 36206763). HPAP is part of a Human Islet Research Network (RRID:SCR_014393) consortium (UC4-DK112217, U01-DK123594, UC4-DK112232, and U01-DK123716). We thank Kerem Fidan for helpful comments on the manuscript draft.

## Supplementary Figures

**Supplementary Figure 1:**
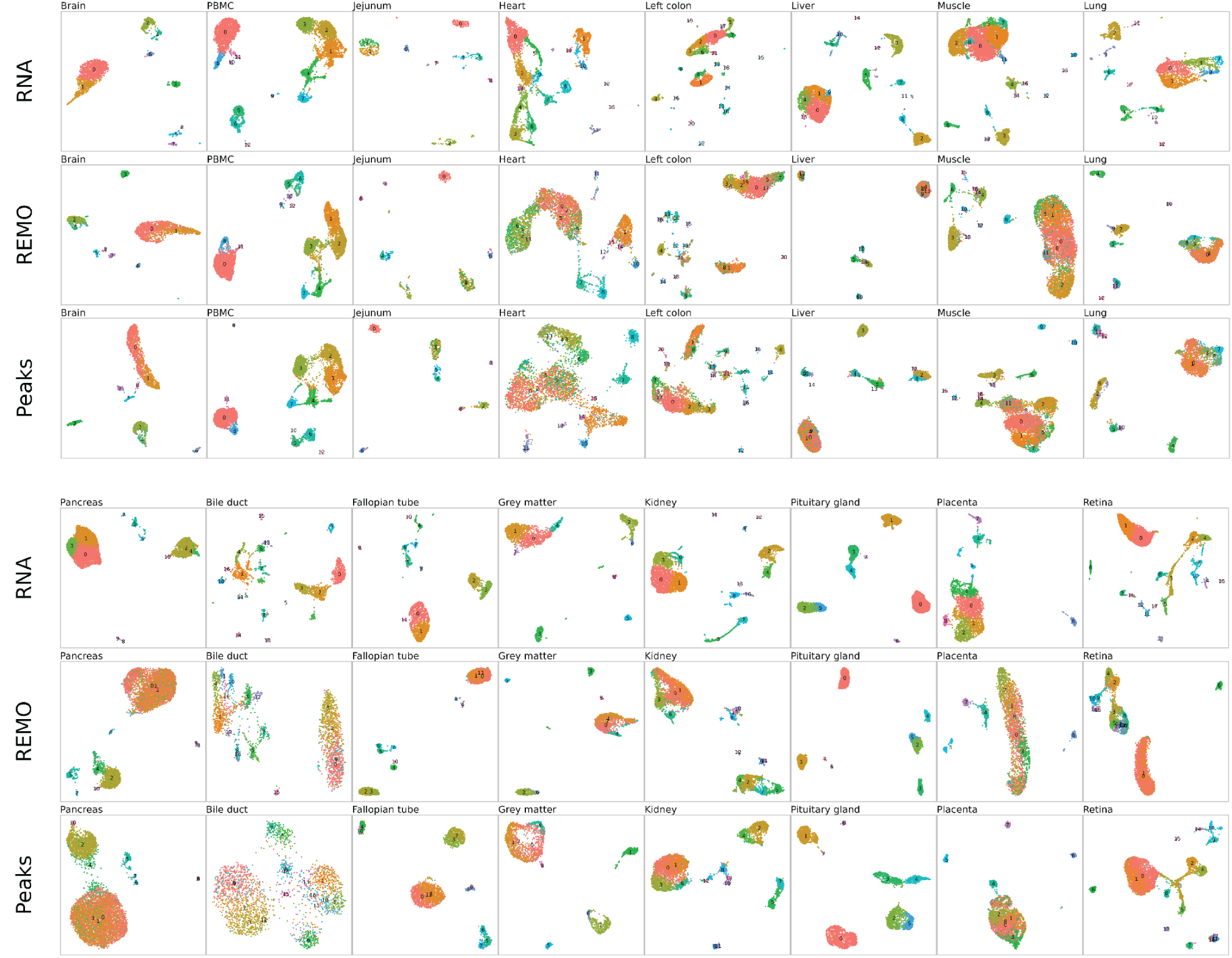
UMAP representations for all multiome datasets. Two-dimensional UMAP representation for 16 multiome datasets processed using gene expression data (RNA) or DNA accessibility data either using REMO or peak calling. For each dataset, cells were annotated by gene expression-based clustering.

